# PPARα-dependent effects in the liver and jejunum of mice in response to pemafibrate

**DOI:** 10.64898/2026.09.30.755633

**Authors:** Frédéric Lasserre, Arnaud Polizzi, Anne Fougerat, Sandrine Ellero-Simatos, Yannick Lippi, Marine Huillet, Céline P Martin, Sarra Smati, Claire Naylies, Géraldine Garcia, Bart Staels, Walter Wahli, Thierry Pineau, Alexandra Montagner, Nicolas Loiseau, Hervé Guillou, Elena Piccinin

**Author notes:** Corresponding authors: Hervé Guillou Elena Piccinin.

## Abstract

Peroxisome proliferator-activated receptor α (PPARα) is a nuclear receptor that plays a central role in metabolic adaptation by regulating gene expression programs involved in lipid utilization and energy homeostasis. Although the hepatic functions of PPARα and its target genes are well characterized, less is known about its transcriptional activity in extrahepatic tissues. Pemafibrate (K-877), a selective PPARα modulator (SPPARMα), is a high-affinity PPARα ligand with improved potency and selectivity compared with conventional fibrates. Here, we investigated pemafibrate-induced tissue-specific transcriptional responses comparing wild type and whole-body PPARα-deficient littermate mice. Twelve-week-old male PPARα+/+ and PPARα-/- mice were treated daily with pemafibrate (0.1 mg/kg) or vehicle by oral gavage for 14 days. We performed genome-wide transcriptomic analyses in the liver, jejunum, kidney, soleus muscle, and BAT. Pemafibrate-induced transcriptional changes were entirely dependent on PPARα, as no differentially expressed genes were detected in PPARα-/- mice. The magnitude and nature of the response were tissue-specific, with the strongest transcriptional effects observed in the liver and jejunum, whereas the kidney, soleus muscle, and BAT showed limited responses. In the liver, pemafibrate primarily induced the expression of genes involved in fatty acid metabolism, mitochondrial function, and energy production. In the jejunum, pemafibrate activated lipid metabolic programs together with tissue-specific pathways related to cellular homeostasis. Collectively, our findings show that pemafibrate is a highly selective PPARα activator with a preferential transcriptional activity in the liver and intestine after oral administration. This study provides a resource for defining tissue-specific PPARα-responsive gene networks and expands our understanding of PPARα functions beyond the liver.

## Introduction

Obesity has reached pandemic proportions, affecting more than two billion adults worldwide and representing a major global health challenge. It is associated with an increased risk of cardiovascular disease, type 2 diabetes, and metabolic dysfunction-associated steatotic liver disease (MASLD), all of which are characterized by profound disturbances in lipid and glucose metabolism [1]. Understanding the molecular mechanisms that coordinate metabolic adaptation across tissues is therefore essential for developing of effective therapeutic strategies.

The nuclear receptor peroxisome proliferator-activated receptor α (PPARα) is a master transcriptional regulator of systemic energy metabolism. PPARα is highly expressed in several metabolically active tissues, including the liver, intestine, kidney, skeletal muscle, and brown adipose tissue (BAT), where its activation by ligands induces transcriptional programs involved in fatty acid uptake, mitochondrial and peroxisomal β-oxidation, ketogenesis, amino acid metabolism, and glucose homeostasis [2-4]. Among these organs, the liver is by far the best-characterized target tissue of PPARα [5]. Decades of genetic and pharmacological studies have established its essential role in the adaptive response to fasting and lipid overload, and have identified a large repertoire of direct hepatic target genes controlling fatty acid oxidation, ketogenesis, lipoprotein metabolism, and hepatokine secretion. Activation of hepatic PPARα also promotes triglyceride clearance and reverse cholesterol transport by inducing genes involved in lipoprotein metabolism, thereby reducing circulating triglycerides and increasing HDL-cholesterol [6]. In contrast, although PPARα is abundantly expressed in several extrahepatic tissues, its physiological functions, direct target genes, and contribution to tissue-specific metabolic regulation remain much less well understood.

Given its central role in lipid homeostasis, PPARα has long been considered an attractive therapeutic target for dyslipidemia and associated metabolic disorders. Fibrates, including fenofibrate and clofibrate, have been used clinically for decades to lower plasma triglycerides through PPARα activation. However, these first-generation agonists display relatively low potency and limited receptor selectivity, resulting in variable efficacy and off-target effects that can restrict their therapeutic benefit [7-9].

The development of pemafibrate (K-877), a selective PPARα modulator (SPPARMα), represents a major advance in PPARα pharmacology. Owing to its unique Y-shaped chemical structure, pemafibrate fits optimally within the Y-shaped ligand-binding pocket of PPARα, inducing receptor conformations that promote selective cofactor recruitment and enhanced transcriptional activity [10, 11]. Compared with conventional fibrates, pemafibrate exhibits substantially greater potency and selectivity for PPARα. . These properties, together with its triglyceride-lowering activity, have generated considerable interest in its therapeutic potential for dyslipidemia and other metabolic diseases, including MASLD [12-16]. However, despite improvements in triglycerides and other lipid parameters, the PROMINENT clinical trial showed no significant reduction in the incidence of cardiovascular events in patients receiving pemafibrate compared with those receiving placebo [17]. These findings highlight the need to better understand the biological effects of pemafibrate to help refine its potential clinical applications.

An important unresolved question is how selective PPARα activation shapes transcriptional responses across different tissues. Most studies have focused on the liver, where PPARα-regulated pathways and target genes have been extensively characterized [18], whereas comparatively little is known about the molecular programs that PPARα controls in other metabolically active organs. Because PPARα activity is determined not only by ligand binding but also by tissue-specific chromatin accessibility, transcription factor networks, and co-regulator availability, activation of the same receptor is likely to produce distinct transcriptional outputs depending on cellular context. A comprehensive understanding of these tissue-specific responses is therefore essential for elucidating both the physiological functions of PPARα and the mechanisms underlying the therapeutic actions of selective PPARα modulators.

In thisstudy, we systematically characterized the transcriptional responses to pemafibrate across multiple metabolically active organs in wild-type and whole-body PPARα-deficient mice. By combining pharmacological activation with genetic ablation of PPARα, we defined the tissue specificity of pemafibrate action and identified distinct PPARα-dependent transcriptional programs in the liver and jejunum, providing new insights into the organ-specific mechanisms underlying selective PPARα modulation.

## Results

### Tissue-specific regulation of gene expression by pemafibrate

To examine the effect of pemafibrate on metabolically active tissues, PPARα^+/+^ and PPARα^-/-^ mice were fed a chow diet and treated daily via oral gavage with 0.1 mg/kg pemafibrate or vehicle for 14 days (Figure 1A). Pemafibrate did not impact total body weight and perigonadal white adipose tissue mass, but it increased liver weight in PPARα^+/+^ mice (Figure 1B). PPARα deletion was confirmed by the absence of expression and induction of three well described PPARα target genes - *Cytochrome p450 4a14* and *4a10 (Cyp4a14* and *Cyp4a10*) and *Enoyl-CoA Hydratase and 3-Hydroxyacyl-CoA Dehydrogenase (Ehhadh)* – in PPARα^-/-^ mice, whose expression was induced in the liver of PPARα^+/+^ mice treated with pemafibrate (Figure 1C).

**Figure 1.**
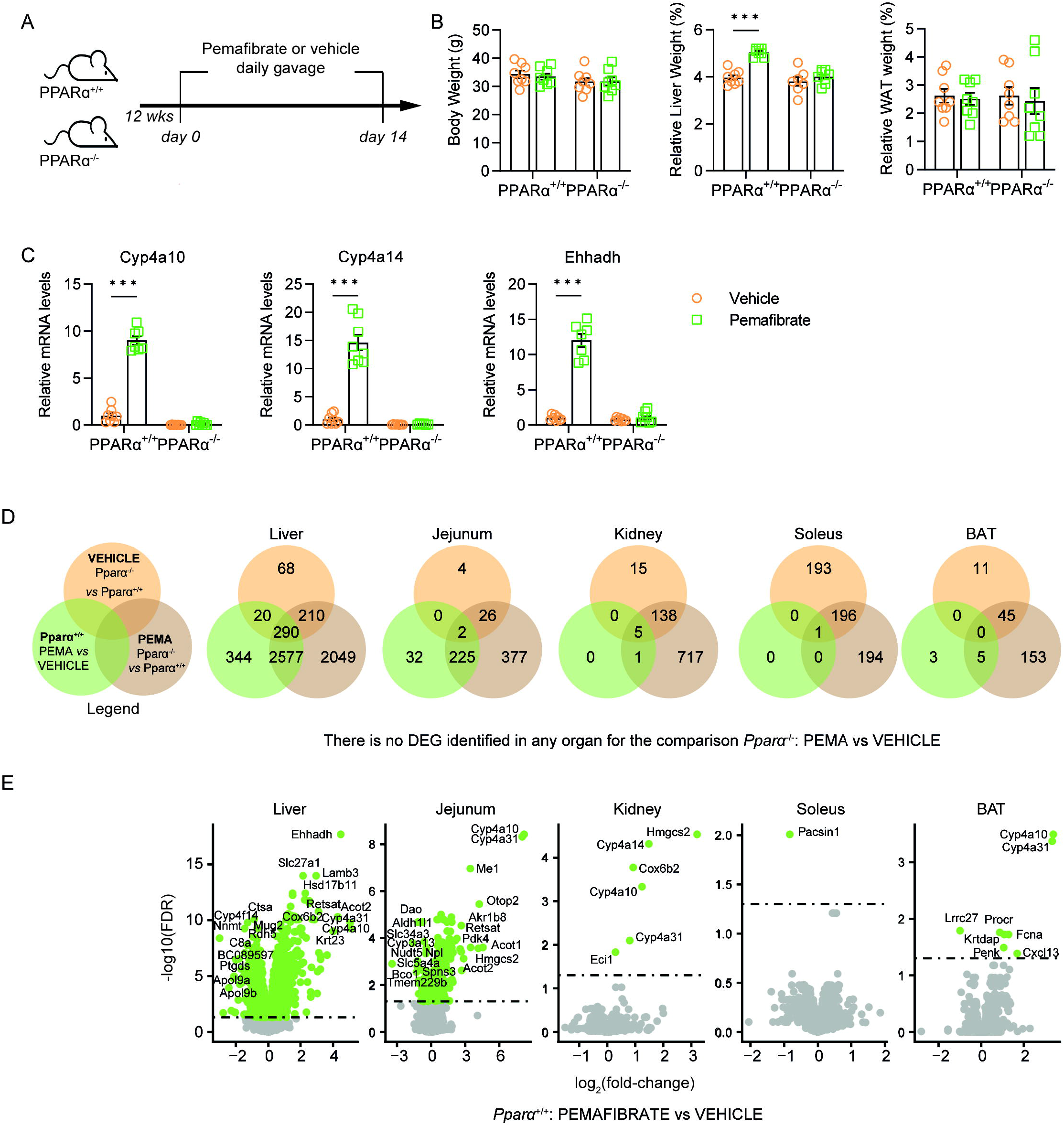
Pemafibrate administration to PPARα^+/+^ and PPARα^-/-^ mice. (A) Experimental design. (B) Body weight, liver weight, and perigonadal white adipose tissue (WAT) mass. (C) Hepatic gene expression of known PPARα target genes. mRNA levels were determined via real-time qPCR, using TATA-binding protein (Tbp) as an endogenous control. Results are expressed as mean±SEM (n=8 mice/group). Comparisons between different groups were performed using 2-Way ANOVA followed by Tukey’s post hoc test (***p < 0.001). (D) Venn diagram analysis of all the differentially expressed genes (FDR < 0.05) in the liver, jejunum, kidney, soleus, and brown adipose tissue (BAT) between PPARα^+/+^ and PPARα^-/-^ mice after treatment with Pemafibrate or vehicle. All comparisons test the treatment effect within the same genotype or the genotype effect in mice receiving the same treatment. (E) Volcano plot showing differential gene expression between Pemafibrate- or vehicle-exposed PPARα^+/+^ mice across the liver, jejunum, kidney, soleus, and BAT. Genes in green are considered differentially expressed (FDR <0.05). The top 10 most up- and down-regulated genes are labeled with scores based on log2FC multiplied by -log10(FDR ) of the Pemafibrate effect in PPARα^+/+^ mice).

We then conducted a genome-wide transcriptomic microarray analysis to characterize the impact of pemafibrate administration on metabolically relevant organs, the liver, jejunum, kidney, soleus and BAT. Pemafibrate was confirmed to be a highly selective PPARα agonist. Consistent with its PPARα-dependent mechanism of action, no differentially expressed genes were detected in any organ of PPARα^™/™^ mice treated with pemafibrate, indicating the absence of detectable PPARα-independent transcriptional effects of the drug. The comparison of differentially expressed genes (DEGs) in each tissue of PPARα^+/+^ mice revealed that pemafibrate has a major effect on the hepatic and intestinal transcriptomes, while its effects are almost null in the kidney, soleus, and BAT (Figure 1D, green circles). The liver exhibited the largest number of modulated genes after pemafibrate administration. Among the 5558 hepatic DEG, 5126 (92%) are genes regulated upon pemafibrate exposure. Of these, 2577 (46%) were shared with the comparison between pemafibrate-treated PPARα^+/+^ and PPARα^-/-^ mice, indicating a specific PPARα-dependent response to pemafibrate (Figure 1D). However, a large number of genes (2049, 37%) were uniquely associated with the genotype effect in pemafibrate-treated mice, suggesting suggesting a potential genotype-dependent response to pemafibrate(Figure 1D, brown circles). Notably, 210 (3.8%) genes responded to PPARα activity independently of pemafibrate treatment, while 290 (5.2%) genes were in the intersection, indicating complex regulation by both PPARα and pemafibrate. Indeed, under physiological conditions, endogenous ligands can activate a basal PPARα-dependent response. Many endogenous fatty acids and lipid-derived metabolites, originating from the diet or synthesized endogenously, are natural PPARα ligands and can activate the receptor with physiologically relevant affinities, thereby sustaining basal transcriptional activity even in vehicle-treated animals [19-21].

The transcriptomic effect of pemafibrate is also particularly relevant in the jejunum, where 630 (95%) genes are modulated by its administration in PPARα^+/+^ mice (Figure 1E). Of these, 225 (36%) genes are shared between those regulated by pemafibrate in PPARα^+/+^ mice and those differentially expressed between pemafibrate-treated PPARα^+/+^ and PPARα^™/™^ mice, indicating that the transcriptional response to pemafibrate is dependent on PPARα. By contrast, 26 (4.1%) genes are shared between genotype comparisons irrespective of the treatment. Unlike the liver, only 2 genesfound in the intersection, and are affected by PPARα expression and pemafibrate treatment. Conversely, in the kidney and soleus, only a few genes are modulated by pemafibrate and PPARα expression (5 and 1 gene, respectively), while no gene was detected in the intersection in BAT (Figure 1E).

Consistently, the volcano plots show that pemafibrate treatment can either increase or decrease the expression of many genes in the liver or jejunum of PPARα^+/+^ mice compared to vehicle administration (Figure 1E). Notably, the overall pattern is similar between the two organs, many genes are significantly upregulated or downregulated, but the specific genes and magnitude of change differ. Conversely, pemafibrate affected only a few genes in the kidney and BAT, and only one gene was significantly downregulated by pemafibrate in the soleus (Figure 1E). As expected, *Cyp4a31* and *Cyp4a10* show the largest changes across the tissues examined, except for the soleus (Figure 1E).

The substantial variation in the number of shared and specific differentially expressed genes between the liver, jejunum, kidney, soleus and BAT emphasises the tissue-specific nature of the transcriptional response to pemafibrate-activated PPARα .

### Hepatic and intestinal-specific biological pathways sensitive to pemafibrate administration

The liver and jejunum were the organs most affected by pemafibrate treatment. Therefore, we submitted the DEGs identified in the liver and jejunum for further analysis to determine the most affected biological processes in these two tissues.

Gene set enrichment analysis (GSEA) comparing treatment effects in the livers of PPARα^+/+^ mice revealed significant involvement of pathways related to mitochondrial function and metabolism. These include mitochondrial translation, aerobic respiration, fatty acid β-oxidation, ATP metabolic processes, and organophosphate metabolism (Figure 2A). Most of these pathways wereupregulated by the treatment (indicated by yellow nodes, positive NES), suggesting that pemafibrate activates mitochondrial and oxidative metabolic processes.

**Figure 2.**
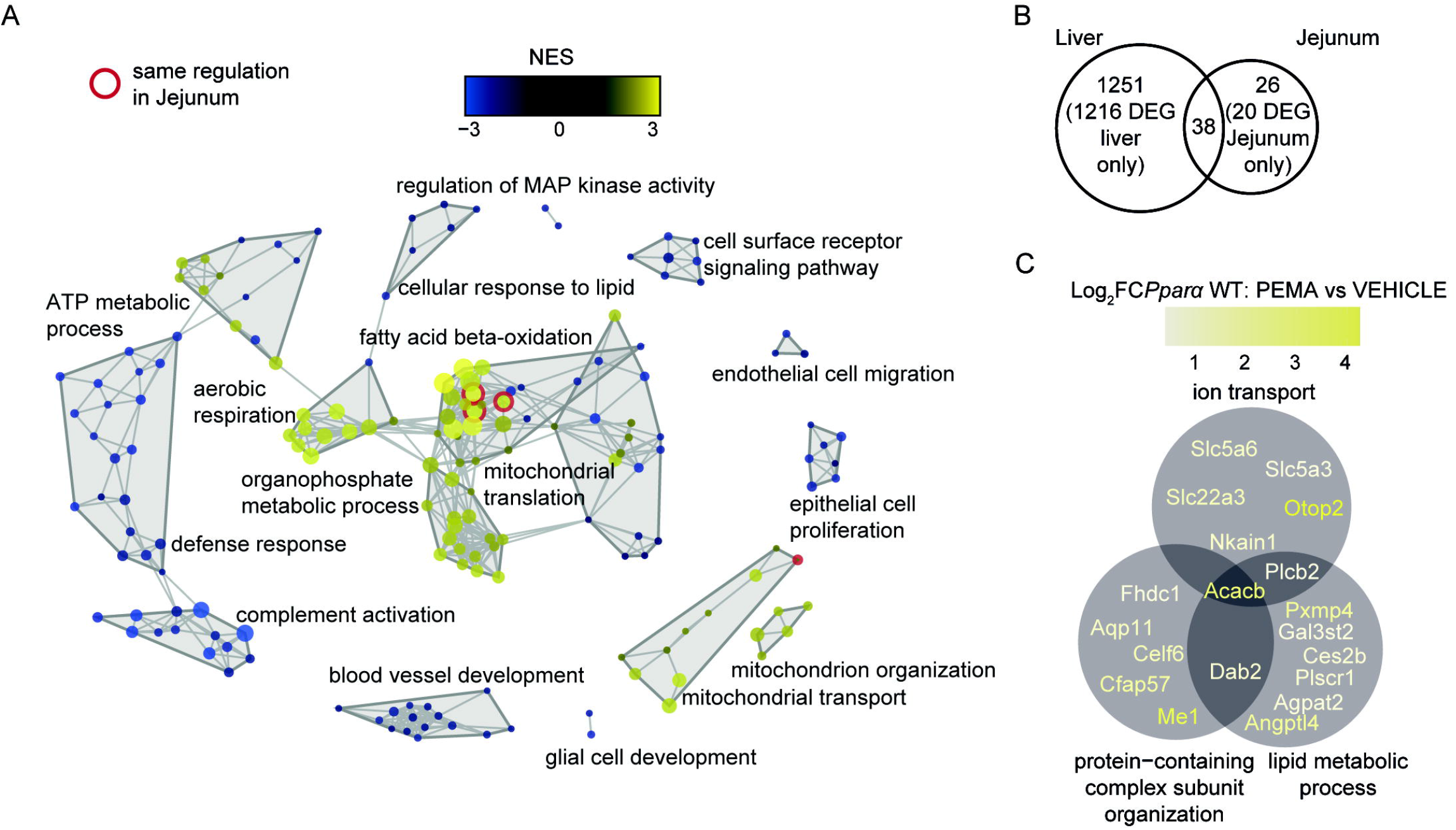
Hepatic and intestinal-specific biological pathways affected by Pemafibrate administration. (A) Network of the most significant categories obtained by testing the Pemafibrate effect in the liver of PPARα^+/+^ mice using GSEA (FDR <0.05, score = signal-to-noise ratio). The color indicates the Normalized Enrichment Score (NES), and the size indicates the -log10 FDR. Categories marked with a red circle are also regulated in the Jejunum in the same direction. (B) Venn diagram showing the number of leading edges from significant categories identified by GSEA (FDR < 0.05) in each organ during Pemafibrate treatment in the liver of PPARα^+/+^ mice. (C) Illustration of the three GO categories significant in GSEA (FDR < 0.05) in the jejunum only, along with their associated leading-edge genes.

While GSEA analysis revealed a strong impact on the hepatic transcriptome identifying 1251 leading-edge genes - specific “core” genes within an enriched pathway that contribute most to the enrichment result – that were specifically regulated by pemafibrate in the liver, only 26 genes were identified as leading in the analysis of the jejunum alone (Figure 2B). The nodes marked in red in Figure 3A represent pathways that were similarly regulated in the liver and jejunum. Although the transcriptional changes in the intestine were modest, they largely aligned with the responses observed in the liver. In fact, a comparison of leading-edge genes between the liver and jejunum showed that pemafibrate modulated 38 genes in both tissues (Figure 2B).

**Figure 3.**
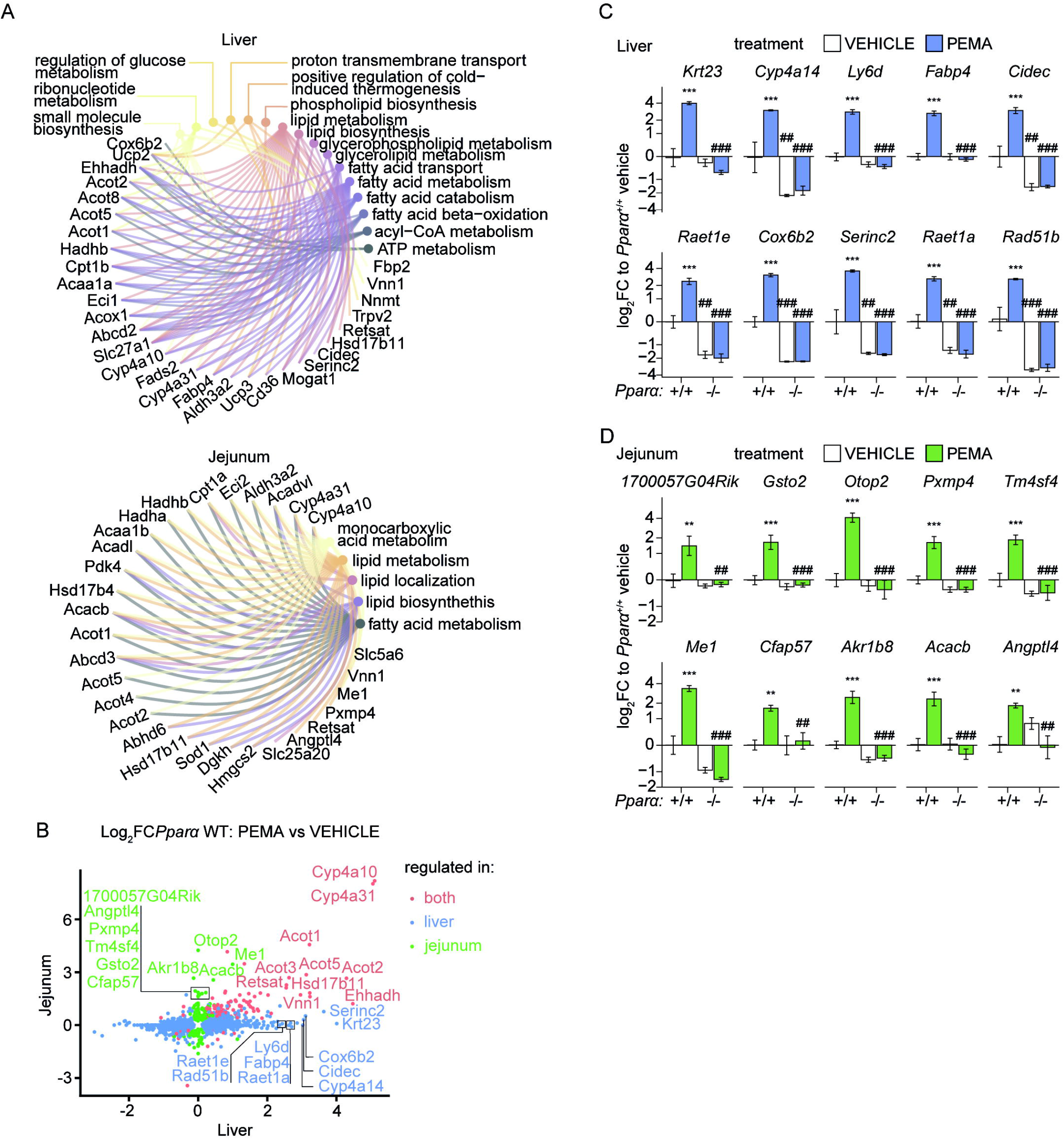
Differentially expressed genes in the liver and jejunum of PPARα^+/+^ mice treated with Pemafibrate. (A) Circular networks of the top 30 most differentially expressed genes per organ (score = log2FC * -log10(FDR) of the Pemafibrate effect in PPARα^+/+^ mice) and their more precise associated GO categories found significant in GSEA (FDR < 0.05). (B) Correlation plot of log2FC of the Pemafibrate effect in PPARα^+/+^ mice in the liver versus the jejunum. Color indicates the significance of the effect in each tissue (FDR<0.05). Bar plots of the top 10 most differentially induced genes specifically in the liver(C) and in the Jejunum (D) (score = log2FC of the Pemafibrate effect in PPARα^+/+^ mice). *: treatment effect; #: genotype effect; * or # FDR <0.05; ** or ## FDR <0.01; *** or ### FDR <0.001. Data are represented as mean ± SEM.

Among the 26 leading-edge genes specifically identified in the jejunum, 73% belong to 3 interconnected GO categories (Figure 2C). One category included genes involved in ion transport, another comprised genes associated with protein-containing complex assembly/organization, and the last included genes related to lipid metabolic processes. These genes were generally upregulated, as expected from the pemafibrate induction of PPARα-dependent genes.

Overall, these data show that pemafibrate robustly activates mitochondria and oxidative metabolic processes in the liver, and induces a smaller but consistent transcriptional response in the jejunum, particularly affecting ion transport and lipid metabolism.

### Differentially expressed genes in the liver and jejunum of PPARα^+/+^ mice treated with pemafibrate

To further characterise the pemafibrate effect in the liver and jejunum of PPARα^+/+^ mice, we selected the top 30 differentially expressed genes to create a network diagram illustrating the relationships between specific genes and metabolic pathways in the two organs.

The pathway-gene network visualisation showed that in the liver, pemafibrate strongly increased the expression of genes involved in fatty acid metabolism and β-oxidation, lipid metabolism, and ribonucleotide metabolism (Figure 3A, top). Key genes in these pathways included *Ehhadh, Cyp4a31, Cyp4a10*, and *Acyl-CoA Thioesterase* 2 (*Acot2)*, consistent with pemafibrate’s known role as a PPARα agonist. Additionally, pemafibrate upregulated proteins involved in fatty acid transport (*Slc27a1, Fabp4*, and *Cpt1b*) and the thermogenesis pathway (*Uncoupling protein 2 and 3, Ucp2* and *Ucp3*).

By contrast, the jejunal response was less extensive but more focused on lipid metabolic pathways and monocarboxylic acid metabolism (Figure 3A, bottom), with key genes such as *Cyp4a10/4a31, Acetyl-CoA carboxylase 2 (Acacb), ATP-binding cassette d3 (Abcd3)*, and *acyl-CoA dehydrogenase very long chain* (*Acadvl)* being affected.

The direct comparison of the transcriptional responses in PPARα^+/+^ mice revealed that most pemafibrate-regulated genes were specific to the liver, while a smaller subset was shared between the liver and the jejunum, and a unique group was specific to the jejunum (Figure 3B). Genes shared between the two organs largely belonged to lipid oxidation, with the highest expression observed for *Cyp4a10* and *Cyp4a31*. Additionally, *Retinol saturase* (*Retstat), Ehhadh, Vanin1 (Vnn1)*, and several *Acot* isoforms are upregulated in both tissues, though to a lesser degree.

We then examined the top 10 DEGs specific to the liver and the jejunum of PPARα^+/+^ mice treated with pemafibrate. The hepatic-specific genes induced by pemafibrate in PPARα^+/+^ but not in PPARα^-/-^ mice are more closely related to lipid metabolism (*Krt23, Lys6, Cidec, Cyp4a14, Fabp4, Rat1e/1a, SerinC2*), with only a few genes involved in mitochondrial (*Cox6b2*) and nucleotide metabolism (*Rad51b*) (Figure 3C). In the jejunum, pemafibrate selectively upregulates genes involved in various cell processes, including lipid metabolism (*Me1, Acacb, Angptl4, Pxmp4, 1700057G04Rik* (recently defined as phospholipid scramblase Plscr1), transport (*Tm4sf4, Otop2*), protein complex (*Gsto2, Akr1b8*), and structural components (*Cfap57*), again exclusively in PPARα^+/+^ mice (Figure 3D).

Overall, the results showed that while both tissues responded to pemafibrate by activating lipid metabolic pathways, the response varied in magnitude. The liver showed a more extensive and stronger activation of a wider array of metabolic and non-metabolic pathways than the jejunum. This further emphasizes the tissue-specific effects of pemafibrate, with the liver displaying the most extensive transcriptional response to the treatment.

## Discussion

PPARα is highly expressed in metabolically active tissues, with the highest levels observed in the liver and brown adipose tissue (BAT), followed by the small intestine, skeletal muscle, and kidney [5, 22]. Based on this expression pattern, we selected these five organs to characterize the pemafibrate-induced tissue-specific transcriptional responses. Importantly, pemafibrate treatment did not induce significant transcriptional changes in PPARα^-/-^ mice in any examined tissues, showing that its *in vivo* activity depends heavily on PPARα and indicating minimal contribution from off-target effects under these experimental conditions.

Pemafibrate administration increased liver weight in PPARα^+/+^ mice, whereas this effect was absent in PPARα-deficient animals. This observation is consistent with previous studies showing that fibrates and other pharmacological PPARα activators induce hepatomegaly and hepatocyte hypertrophy in rodents [23, 24]. However, this proliferative response appears to be largely restricted to rodents and is thought to result from species-specific differences in PPARα structure, expression, and transcriptional activity between mice and humans [25, 26].

Our transcriptomic analysis revealed a striking tissue specificity in the response to pemafibrate. While numerous PPARα-dependent genes were regulated in the liver and jejunum, only a limited number of transcriptional changes were detected in BAT, skeletal muscle, and kidney. This preferential activity in the liver and intestine may be related, at least in part, to the pharmacokinetic properties of pemafibrate. Unlike conventional fibrates such as fenofibrate, which undergo substantial renal excretion and consequently activate renal PPARα signaling [24], pemafibrate is primarily metabolized by the liver and excreted through the bile, with minimal urinary excretion of the unchanged compound [27]. This distribution profile likely contributes to the strong transcriptional response observed in the liver and jejunum. However, whether intestinal PPARα activation results from direct exposure to pemafibrate, biliary delivery of the parent compound, or active metabolites remains unclear.

Across all analyzed organs except soleus muscle, we observed consistent induction of *Cyp4a10* and *Cyp4a31* expression. These genes encode cytochrome P450 enzymes involved in fatty acid ω-hydroxylation and are well-established as PPARα-responsive genes [28-30]. Their conserved regulation across tissues suggests that certain core components of PPARα transcriptional activity are maintained throughout the organism, whereas additional gene programs appear to be strongly influenced by tissue context.

In the liver, pemafibrate predominantly stimulated transcriptional pathways involved in lipid metabolism, particularly fatty acid degradation. Gene set enrichment analysis revealed a strong enrichment of mitochondrialpathways, including mitochondrial translation, aerobic respiration, and ATP metabolic processes in treated PPARα^+/+^ mice (Figure 2A). Consistent with this observation, pemafibrate selectively induced *Cox6b2* expression in the liver, which encode a component of mitochondrial complex IV. Activation of mitochondrial pathways aligns with PPARα’s central role in promoting fatty acid β-oxidation, which generates acetyl-CoA substrates required for tricarboxylic acid cycle activity, oxidative phosphorylation, and ATP production.

Among the genes preferentially induced in the liver, *Lys6, Cidec*, and *Krt23* have previously been associated with lipid accumulation and metabolic liver disease. However, their induction occurred alongside broad activation of lipid catabolic pathways, which likely counterbalances any potential lipogenic or stress-associated effects. Indeed, extensive genetic and pharmacological evidence demonstrates that hepatic PPARα activation protects against excessive lipid accumulation by promoting fatty acid oxidation and maintaining hepatic lipid homeostasis [30-32]. Preclinical and clinical studies consistently indicate that pemafibrate improves several features associated with MASLD, including hepatic steatosis and metabolic dysfunction [33-36]. Moreover, hepatic PPARα expression in humans is inversely associated with metabolic dysfunction-associated steatohepatitis (MASH) severity, and evidence shows that histological improvement correlates with increased PPARα and downstream target gene expression [37], further supporting pemafibrate as a potential therapeutic option for MASH. In the jejunum, pemafibrate strongly activated lipid metabolic pathways, with fatty acid β-oxidation among the most enriched. The most highly regulated genes included *Cpt1a*, encoding the rate-limiting enzyme controlling mitochondrial fatty acid entry, and several *Acot* family members involved in regulating fatty acyl-CoA availability for oxidation. These findings are consistent with previous studies in human jejunal biopsies, in which treatment with a PPARα agonist induced *Cpt1a* and *Acox1 expression*, similarly to what we observed in our treated PPARα^+/+^ mice [38]. Interestingly, *Fabp1*, previously reported as an intestinal PPARα target in obese individuals, was not induced in our model, suggesting that intestinal PPARα regulation depends heavily on physiological and pathological contexts [39, 40]. Intestinal PPARα activity has been implicated in maintaining epithelial adaptation, regulating cell proliferation, differentiation, and apoptosis in the small intestine, influencing villus morphology and absorptive capacity, processes that require substantial metabolic support due to continuous epithelial renewal [41, 42]. Furthermore, several jejunum-specific pemafibrate-responsive genes, including *Gsto2, Pxmp4, Akr1b8*, and *Me1*, are involved in cellular detoxification and redox homeostasis. Intriguingly,redox metabolism was previously marked as upregulated also in the jejunum of mice treated with a selective PPARα agonist WY14643 [41], suggesting that intestinal PPARα contributes not only to lipid utilization but also to maintaining epithelial integrity.

Together, these findings identify the intestine as an important target tissue for pemafibrate action. Given the growing recognition of the gut–liver axis as a key regulator of metabolic liver diseases [40, 43], the contribution of intestinal PPARα activation to pemafibrate’s overall therapeutic effects warrants further investigation.

Although the PROMINENT trial demonstrated that pemafibrate efficiently improves dyslipidaemia, it did not significantly reduce cardiovascular events in patients with type 2 diabetes and mild-to-moderate hypertriglyceridemia [17]. Nevertheless, pemafibrate remains of considerable interest for metabolic liver diseases, where modulation of hepatic lipid metabolism may provide therapeutic benefits [44, 45]. Our findings provide mechanistic insight into the pemafibrate’s tissue selectivity by demonstrating preferential activation of PPARα-dependent transcriptional programs in the liver and intestine, whit limited effects in the kidney and other peripheral tissues.

Beyond these mechanistic insights, our study offers a resource for the scientific community interested in PPARα biology. Although hepatic PPARα signaling and its associated transcriptional targets are well characterized, comparatively little is known about PPARα-responsive gene networks in extrahepatic tissues. By integrating pemafibrate pharmacological activation with PPARα genetic deletion across multiple organs, our dataset identifies tissue-specific PPARα-dependent and PPARα-sensitive genes and provides a framework for exploring the PPARα’s physiological functions beyond the liver.

This study also has limitations. First, experiments were performed exclusively in male mice. Given the well-established sexual dimorphism in lipid metabolism and PPARα signaling [46], future studies that include female cohorts will be important to determine whether the identified transcriptional programs are conserved across sexes. Second, pemafibrate exposure was limited to a relatively short treatment period. Although this approach allowed us to define early transcriptional responses, longer-term studies are needed to evaluate chronic metabolic adaptations and the sustained impact of tissue-specific PPARα activation. Finally, our transcriptomic analyses were based on microarray profiling of whole tissues, while single-cell approaches would be more informative, providing a more detailed understanding of the contribution of individual cell populations to the pemafibrate response. In conclusion, our findings demonstrate that pemafibrate is a highly selective PPARα modulator with a tissue-specific distribution profile. By preferentially activating PPARα-dependent metabolic programs in the liver and intestine while limiting effects in other organs, pemafibrate exhibits a differentiated pharmacological profile compared with traditional fibrates. These findings expand our understanding of tissue-specific PPARα biology and provide a valuable resource for identifying PPARα-responsive pathways involved in metabolic regulation and disease.

## Methods

### Animal experiments

Twelve-week-old PPARα^+/+^ and PPARα^-/-^ male mice were treated daily with a gavage of Pemafibrate (0.1mg/kg) at ZT16 for 14 days. Carboxyl methyl cellulose (0.02mg/mL) was used as a vehicle. Mice were randomly assigned to treatment groups for in vivo studies (n = 4 mice per cage).

At sacrifice, tissues were removed, weighed, snap-frozen in liquid nitrogen, and then stored at -80°C until use.

Mice were housed in a pathogen-free facility at 22±2°C with a 12-h light/dark cycle and had free access to food and water. All procedures complied with the European Union Guidelines for the Use and Care of Laboratory Animals. The experiments followed the ethical protocol approved by the independent ethics committee CEEA-86 Toxcométhique (authorization number 17430-2018110611093660).

### RNA extraction and gene expression

Total RNA was extracted from various tissues using TRI Reagent solution (Thermofisher Scientific). RNA concentration was measured spectrophotometrically (Nanophotometer N60, Implen). RNA was reverse transcribed using the High-Capacity cDNA Reverse Transcription Kit (Thermofisher Scientific) for real-time quantitative PCR (qPCR) analyses. We carried out amplifications on an AriaMx 1.8 Real-Time PCR System (Agilent). qPCR data were analysed with LinRegPCR (v2021.1), and normalised to the levels of TATA-box binding protein (*Tbp*) mRNA.

For microarray analysis, gene expression profiles were obtained at the GeT-TRiX facility (GénoToul, Génopole Toulouse Midi-Pyrénées) using Agilent Sureprint G3 Mouse GE v2 microarrays (8×60K, design 074809) according to the manufacturer’s instructions. For each sample, Cyanine-3 (Cy3) labelled cRNA was prepared from 200 ng of total RNA using the One-Colour Quick Amp Labelling kit (Agilent Technologies) according to the manufacturer’s instructions, followed by purification with Agencourt RNAClean XP (Agencourt Bioscience Corporation, Beverly, Massachusetts). Dye incorporation and cRNA yield were checked using Dropsense™ 96 UV/VIS droplet reader (Trinean, Belgium). 600 ng of Cy3-labelled cRNA were hybridised to the microarray slides following the manufacturer’s instructions. Immediately after washing, the slides were scanned with an Agilent G2505C Microarray Scanner using Agilent Scan Control A.8.5.1 software, and fluorescence signals were extracted using Agilent Feature Extraction software v10.10.1.1 with default parameters.

Microarray data and experimental details are available in NCBI’s Gene Expression Omnibus [47], and can be accessed through GEO Series accession number GSE339189(https://www.ncbi.nlm.nih.gov/geo/query/acc.cgi?acc=GSE339189).

### Bioinformatic data analysis

We analysed microarray data using R (R Core Team, 2018) and Bioconductor packages as described in GEO accession GSE339189. Raw data (median signal intensity) were filtered, log2 transformed, and normalised using the quantile method [48].

We fitted a model using the *limma* lmFit function [49]. We performed pairwise comparisons between biological conditions using specific contrasts. A correction for multiple testing was applied using the Benjamini-Hochberg procedure to control the False Discovery Rate (FDR). Probes with FDR ≤ 0.05 were considered to be differentially expressed between conditions.

We conductedGene Set Enrichment Analysis (GSEA) using *fgsea* v1.34.2 [50]. We retrieved Gene Ontology (GO) annotations from *org*.*Mm*.*eg*.*db* v3.21.0 (Carlson M, 2025, doi:10.18129/B9.bioc.org.Mm.eg.db). All terms linked to each gene where retained for the analysis. The GSEA score was the signal-to-noise ratio (SNR = (µ_1_-µ_2_)/(σ_1_+σ_2_)) for each pair of tested comparisons arising from differential analysis, where µ is the mean of a group and σ is its standard deviation). Networks were calculated with *igraph* v2.2.1 [51] and plotted with *ggraph* v2.2.2 [52]. A semantic similarity matrix was computed for the GO category network using *GoSemSim* [53], and a cutoff of 0.45 to Wang’s distance was used to determine the edges. Venn diagrams were plotted using *venn* v1.12 [54]. Volcano plots, correlation plot and barplots from microarray experiments were genrated using *tidyverse* [55]. All R scripts and processed data are available at [56].

## Statistical analysis

All results are presented as mean ± SEM. Grubbs’ test was used to identify and exclude outliers from the analyses. Statistically significant differences between the four groups were determined using the 2-way ANOVA followed by Tukey’s post hoc test. Differences are considered statistically significant with P values lower than 0.05 (*p < 0.05, **p < 0.01, ***p < 0.001). We performed statistical analyses using Prism 9 software (GraphPad Software Inc.).

## Author contributions

Design of the work: FL, NL, HG, EP

Acquisition, analysis, and interpretation of data: FL, AP, AF, SES, YL, MH, CPM, CN, GG, TP, EP

Drafting the work: FL, AP, HG, EP Reviewing the work: SS, BS, WW, AM

## Acknowledgements

We thank all members of the EZOP staff, the GeT-Trix Genotoul facility for their help. This work was supported by Agence Nationale de la Recherche (ANR Hepatologic ANR-21-CE14-0079-01 to HG) and Fondation pour la Recherche Médicale (Equipe FRM EQU202303016327 to HG).

## Abbreviations

Abcd3: ATP-binding cassette d3
Acacb: Acetyl-CoA carboxylase 2
Acadvl: acyl-CoA dehydrogenase very long chain
Acot: Acyl-CoA Thioesterase
BAT: brown adipose tissue
Cyp4a10: Cytochrome p450 4a10
Cyp4a14: Cytochrome p450 4a14
DEG: differentially expressed genes
Ehhadh: Enoyl-CoA Hydratase and 3-Hydroxyacyl-CoA Dehydrogenase
GSEA: Gene set enrichment analysis
MASLD: metabolic dysfunction-associated steatotic liver disease
PPARα: Peroxisome proliferator-activated receptor α
Retstat: Retinol saturase
Ucp: Uncoupling protein
Vnn1: Vanin1

## Notes

*Conflict of Interest:* none

### Competing Interest Statement

The authors have declared no competing interest.

